# The role of ecosystem engineers in shaping the diversity and function of arid soil bacterial communities

**DOI:** 10.1101/2020.12.15.422998

**Authors:** Capucine Baubin, Arielle M. Farrell, Adam Šťovíček, Lusine Ghazaryan, Itamar Giladi, Osnat Gillor

## Abstract

Ecosystem engineers (EEs) are present in every environment and are known to strongly influence ecological processes and thus shape the distribution of species and resources. In this study, we assessed the direct and indirect effect of two EEs (perennial shrubs and ant nests), individually and combined, on the composition and function of arid soil bacterial communities. To that end, top soil samples were collected in the Negev Desert Highlands during the dry season from four patch types: (1) barren soil; (2) under shrubs; (3) near ant nests; or (4) near ant nests situated under shrubs. The bacterial composition was evaluated in the soil samples (fourteen replicates per patch type) using 16S rRNA gene amplicon sequencing, together with physico-chemical measures of the soil, and the potential functions of the community. We have found that the EEs differently affected the community composition. Indeed, barren patches supported a soil microbiome, dominated by *Rubrobacter* and *Proteobacteria*, while in EE patches the *Deinococcus-Thermus* phylum was dominating. The presence of the EEs similarly enhanced the abundance of phototrophic, nitrogen cycle and stress- related genes. In addition, only when both EEs were combined, were the soil characteristics altered. Our results imply that arid landscapes foster unique communities selected by each EE(s), solo or in combination, yet these communities have similar potential biological traits to persist under the harsh arid conditions. Environments with multiple EEs are complicated to study due to the possibility of non-additive effects of EEs and thus further research should be done.

**IMPORTANCE:** Ecosystem engineers are organisms that can create, modify, or maintain their habitat. They are present in various environments but are particularly conspicuous in desert ecosystems, where their presence is tightly linked to vital resources like water or nutrients. Despite their key role in structuring and controlling desert ecosystems, joint engineering, and their effect on soil function, are unknown. Our study explores the contributions of key ecosystem engineers to the diversity and function of their soil microbiome allowing better understanding of their role in shaping habitats and engineering their activity

## INTRODUCTION

Hot desert environments are characterized by long droughts interspersed by intermittent and unpredictable rain events. Water and nutrients in hot desert environments are scarce and unevenly distributed across the land, resulting in patches of contrasting productivities. High-productivity patches, also called resource islands, are defined by large concentrations of organic matter and nutrients (1–3). These resource islands can be formed through the redistribution of nutrients and water by ecosystem engineers (EEs), such as perennials or invertebrates (4, 5). EEs are also known for impacting many parts of the environment, such as soil features, or community composition of microorganisms (6).

An EE is an organism that, directly or indirectly, modifies the availability of resources to other organisms by transforming the physical state of abiotic and/or biotic components of the ecosystem, *sensu* Jones et al. (1994). The impacts of EEs range from physical, through the creation of biogenic structures (e.g. tunnels) (9); to chemical, through the production of compounds that have physiological effects (e.g. root exudates) (10)(10); to biological, through organisms behaviour (e.g. seed dispersal) (11). In drylands, resources, such as nutrients or water, are often concentrated around EEs, boosting the development of diverse populations of annuals and invertebrates (12), as well as microbial communities, including bacteria (13–15), fungi (16), and protists (17).

In desert ecosystems, ants are a notable example of EEs (14). They redistribute resources by bringing soil from the deep layers to the upper layers (bioturbation) and by gathering, storing, and ejecting food items, such as plant material, or dead invertebrates in and around the nest (18, 19). EEs in arid environments also include perennial shrubs (20–24). Their root systems create a soil mound that traps litter, and seeds, allowing for higher water infiltration. The root exudates increase the content of organic matter and the shrub canopies decrease evaporation, leading to prolonged water availability following a rain event (13)(13). The presence of shrubs alters the course of water run-off (6), which impacts the locations of available water for soil microbial communities. In addition, root systems have their own microbiome, which collaborates with the soil microbial community (25).

The role of both ants and perennial shrubs as EEs were reported in various environments (2, 7, 26–29). However, we know little about their joint effect in arid ecosystems. We hypothesized that each EE would shape the soil bacterial communities differently via changes in the soil physico-chemical properties. We further predicted that the effects of shrub canopy and ant nest are non-additive and thus their combination cannot be predicted by the separate effects of each EE. To test our hypotheses, we explored arid soil bacterial microbiomes and soil chemical features during the dry season of 2015. We sampled four different patches: under *Hammada scoparia* shrubs; near the nest openings of the harvester ant, *Messor ebeninus*; in combined patches of nests under shrubs; and in barren soil.

## MATERIALS AND METHODS

### Sampling

The study was conducted in a long-term ecological research (LTER) site in the Central Negev Desert, Israel (Zin Plateau, 34°80′E, 30°86′N). It is characterised by a 90 mm annual rainfall and average monthly temperatures fluctuating from 13°C (January) to 35°C (August). Vegetation is scarce and dominated by the perennial shrubs *Hammada scoparia* and *Atriplex halimus* (30).

Sampling was conducted as previously described (Baubin et al., 2019) with slight modifications, such as the inclusion of Shrub&Nest samples. To summarize, we sampled four distinct patch types: (1) barren soil (Barren); (2) under the canopy of *H. scoparia* (Shrub); (3) 20-30 cm from the main opening of the nest of *M. ebeninus* (Nest); and (4) 20-30 cm from ant nest’s opening that was situated under a shrub canopy (Shrub&Nest). Samples were collected in October 2015, after an eight-month drought.

We sampled 14 random experimental blocks, from each of the four patches (4 patch types x 14 blocks = 56 samples). All samples were collected from the top 5 cm of the soil after removing crust and debris, then processed within 24 hours of collection. In the lab, the soil from two adjacent blocks was composited resulting in 28 samples that were further processed. Each sample was sieve-homogenized through a 2 mm mesh. 5 g of soil were stored in −80°C for molecular analysis, 20 g were used for water content analysis and the rest was dried at 65°C and used for physico-chemical analysis.

### DNA extraction, amplification, and sequencing

Total nucleic acids were extracted from 0.5 g of soil as previously described (31), purified with the ExgeneTM Soil SV kit (GeneAll, Seoul, S. Korea) according to the manufacturer’s instructions. The 16S rRNA encoding genes V3-V4 region was amplified using 341F and 806R primer (32). The PCR reaction consisted of 2.5 μl 10x standard buffer, 10 μM primers, 0.8 mM dNTPs, 0.4 μl DreamTaq DNA polymerase, 4 μl template, 1 mM bovine serum albumin (Takara, Kusatsu, Japan) and 12.6 μl Milli-Q water. Triplicate PCR reactions (95°C for 30 secs; 28 cycles of 95°C for 15 secs, 50°C for 30 secs, 68°C for 30 secs; 68°C for 5 min) were pooled and amplicon concentration and purity were determined by electrophoresis, Nanodrop (ND-1000, Thermo Fisher Scientific, Waltham, MA, USA). The amplicon libraries were constructed and sequenced on the Illumina MiSeq platform (2×250 pair-end) at the Research Resources Centre at the University of Illinois.

### Soil physico-chemical analysis

The physico-chemical parameters of the soil samples were assessed following the standard methods (33). Water content was measured by gravimetry. Other parameters were measured as follows by the Gilat Hasade Services Laboratory (Moshav Gilat, Israel): organic matter (OM) content by dichromate oxidation; nitrate (NO_3_^−^) through aqueous extract; ammonium (NH_4_^+^) through KCl solution extract; phosphorus (P) by sodium bicarbonate extract; and pH in saturated soil extract. The soil parameters were plotted using a Principal Component Analysis (PCA) (stats package (34)) and the significance of difference between patches was evaluated using a non-parametric test: Kruskal-Wallis test and a post-hoc Dunn test (35–37).

### Community analysis

The results were analysed using QIIME2 (38) and Dada2 (39), following the NeatSeq-Flow pipeline. The taxonomic assignment was done using Silva (version 132) (40), through QIIME2 and the statistical analysis was done using R (34). A NMDS plot was created using the Bray-Curtis dissimilarity and the significance of differences between patch types was analysed using ANOSIM (vegan package (41)). The taxonomy was plotted using a stacked bar plot and the significance of difference between patch types was assessed using a non-parametric test: Kruskal-Wallis test and a post-hoc Dunn test (35–37). All sequences retrieved in this study were uploaded to BioProject (https://www.ncbi.nlm.nih.gov/bioproject) under the submission number PRJNA484096.

### Functional Prediction

The prediction of function of the 16S amplicons was done with Piphillin using the KEGG database (October 2018). Piphillin generates a genome abundance table that is normalized to the 16S rRNA copy number for each genome (42, 43). To analyse the arid soil microbial functionality, we selected metabolisms and respective genes related to arid soil using groups and genes from the KEGG database (44). We selected steps in metabolic pathways for different methods of harvesting energy (organotrophy, lithotrophy and phototrophy) (45–48), for parts of the nitrogen cycle (49), and for the survival of the individual during a drought (DNA conservation and repair, sporulation and Reactive Oxygen Species (ROS)-damage prevention) (50–57). Then, we looked for each step in the KEGG database and picked out genes of interest to build our own database. The assignment of function to the KEGG numbers was done in R. The significance of the differences between patch types in predicted functionalities was evaluated using a non-parametric test: Kruskal-Wallis test and a post-hoc Dunn test (35–37) and boxplots were created in R.

## RESULTS

### Soil physico-chemical characteristics

The PCA (Figure 1) tends to show a difference in the soil characteristics (listed in Table S4) between the Shrub&Nest and the other patches (barren, and with one EE: barren, nest, and shrub). Therefore, we will present the average of these other patches compared to the Shrub&Nest average. This difference is primarily explained (up to 77.2% and 22.4%) by the high concentrations of NO_3_ (4.7 mg/kg compared to 30 mg/kg, respectively) and P (22 mg/kg compared to 54 mg/kg, respectively). When verifying with a Kruskal-Wallis test and a Dunn test on the values of these soil variables (Table S5), we see that the differences between patch types are significant (Shrub&Nest vs all other patches, p < 0.05). Patches with two EE also have a significantly higher concentration of NH_4_ (9.72 mg/kg) and OM (8.21%) compared to all other patches (NH_4_ mean: 5.62 mg/kg, p-value <0.05; OM mean: 5.51%, p ≤ 0.05). The water content and pH do not show significant differences between patches (Table S5).

**Figure 1.**
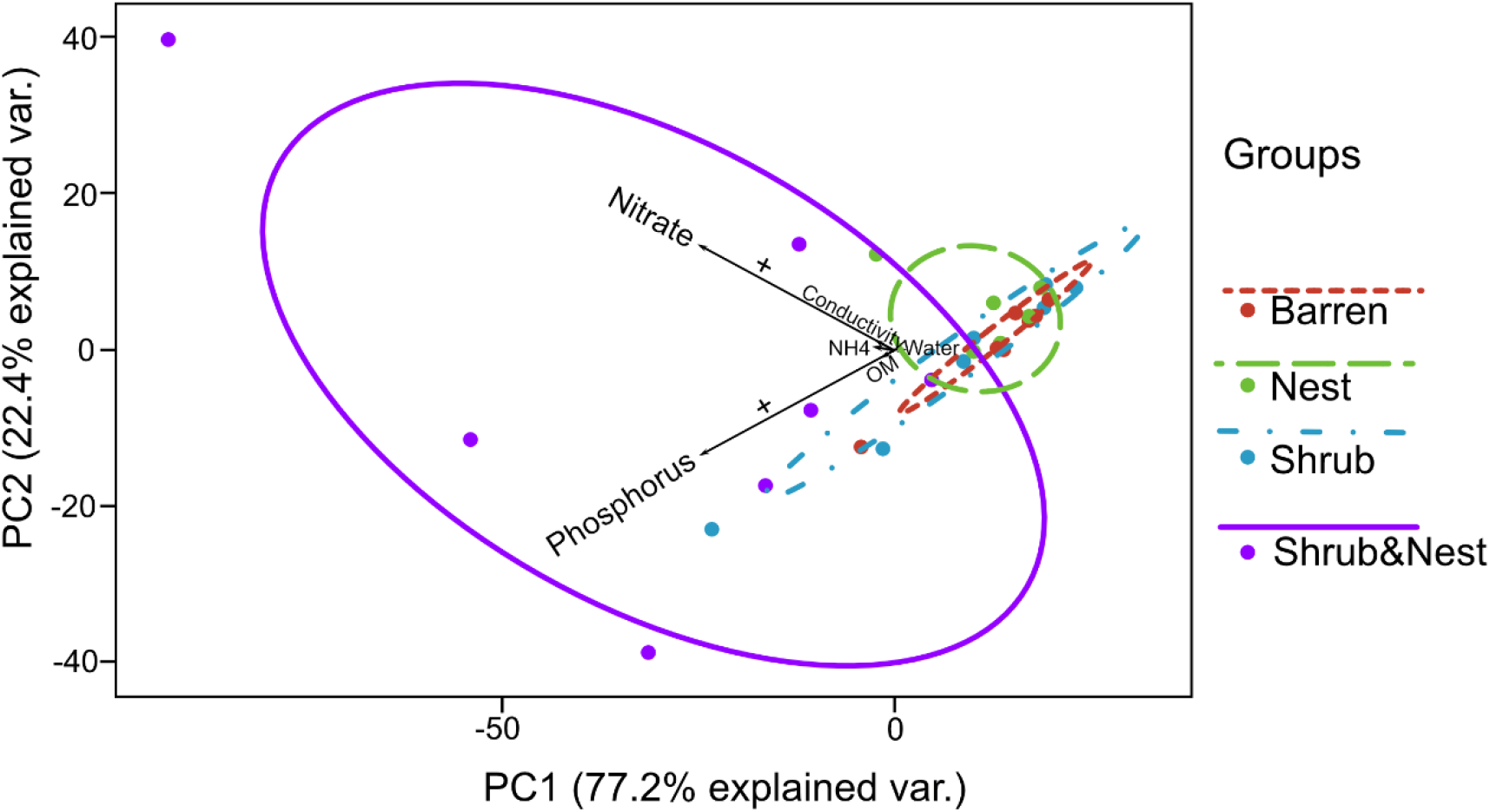
Principal Component Analysis of the soil parameters (Nitrate, Phosphorus, Conductivity, NH4 = Ammonium, OM = Organic Matter content, Water = Water content). The plus signs on the nitrate and the phosphorus vector show an increase in concentration in the Shrub&Nest patches.

### Taxonomy

The summary of the sequence analysis can be found in Table S1. DADA2 analysis yielded 2318 ASVs and the NMDS results (Figure 2) show that there are significant differences in microbial community between patch types (ANOSIM, R= 0.28247; p = 0.001). Most notably, the microbial communities that were sampled in barren soil patches showed high similarities between blocks and were different from those of other patch types (high clustering of barren soil sampling points in the NMDS space). A similar pattern was found for the nest patch type. In contrast, the dissimilarities in community composition within the patch types that included shrubs (Shrub and Shrub&Nest) were high (large scatter of sampling points in the NMDS space).

**Figure 2.**
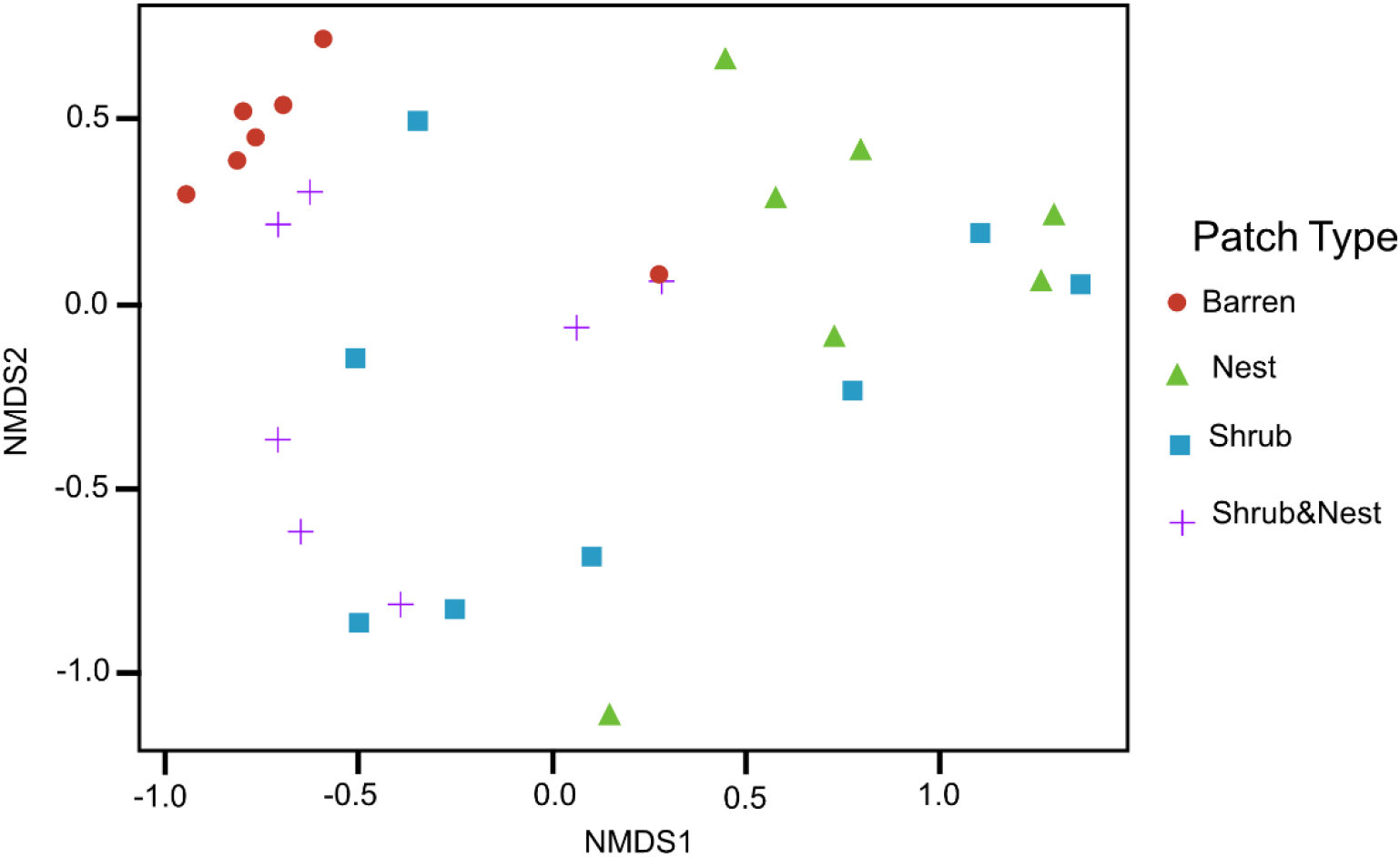
Non-Metric Multidimensional Scaling (NMDS) of the soil 16S microbial communities in the dry season under different patch types. The patch types are significantly different from each other (ANOSIM, R= 0.28247; p-value = 0.001)

The community was mostly composed of *Actinobacteria, Proteobacteria, Deinococcus-Thermus, Bacteroidetes* and *Firmicutes* (Figure 3). The relative abundance for each phylum is detailed in Table S2. We focused on the results of the main three phyla: *Actinobacteria, Deinococcus-Thermus* and *Proteobacteria*. Using pair-wise comparisons, we saw that shrub patches and nest patches had similar communities (no significant differences, p > 0.05) and, therefore, could be considered as patches with a single EE. For these, an average relative abundance of nest and shrub patches is presented. In the *Actinobacteria* phylum, patches with one EE had significantly lower relative abundance than barren patches (one EE: 9 % vs barren patch: 35% p < 0.005) or patches with two EEs (17%, p-value: 0.02). For the *Deinococcus-Thermus* phylum, barren patches had significantly lower relative abundance than patches with one or two EEs (Barren: 3%; vs one EE : 25%; vs two EEs: 9%, p < 0.05). A similar pattern was detected in the *Proteobacteria* phylum (Barren: 38%; vs one EE: 44%; vs two EEs: 39%, p < 0.05). All p-values can be found in Table S3.

**Figure 3.**
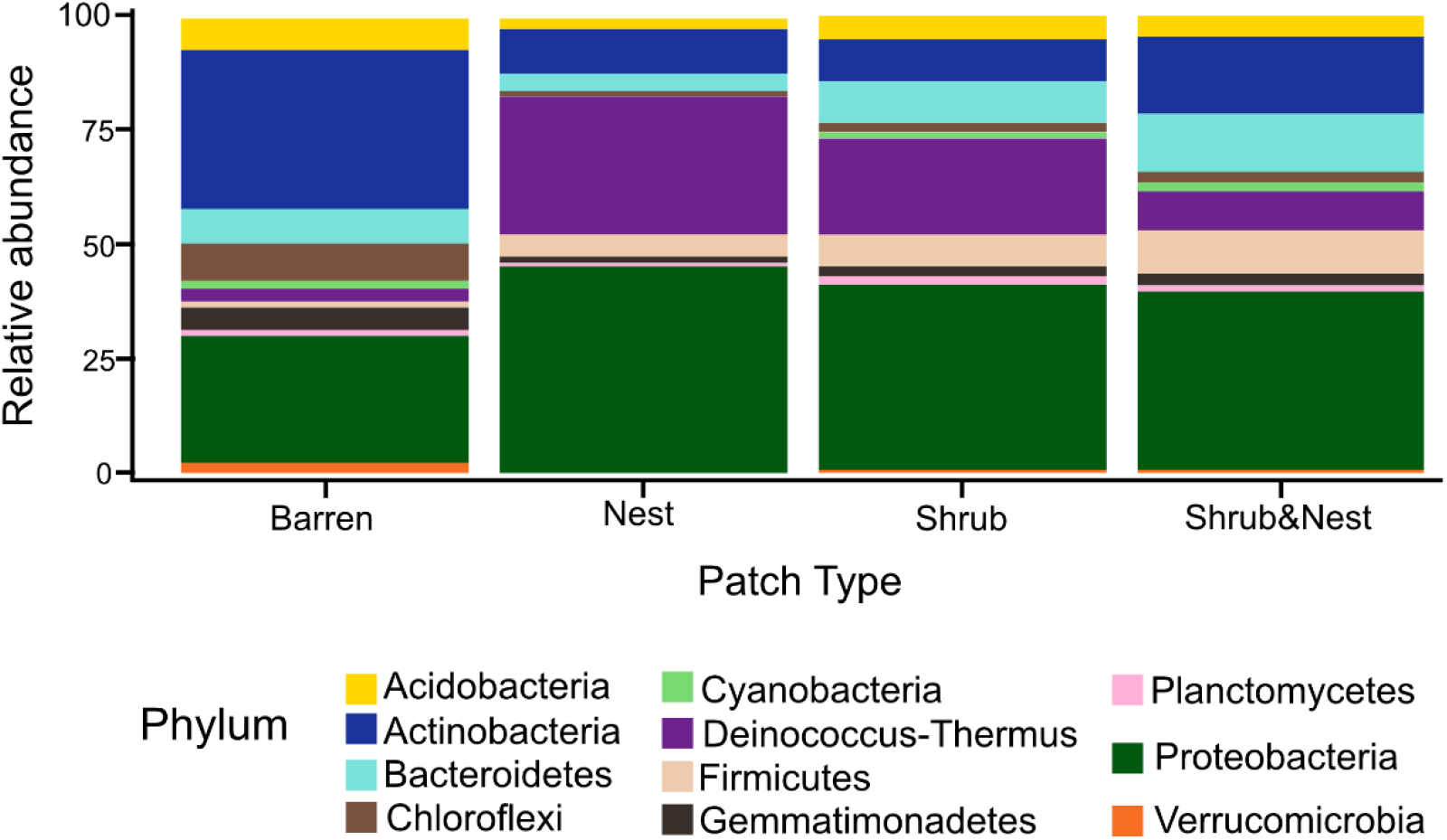
Barplot of the relative abundance (in %) of the most abundant phyla in the soil microbial community in the dry season under different patch types (phyla with a relative abundance > 0.05%). The relative abundance of *Deinococcus-Thermus* increases when one EE is present while the population of *Actinobacteria* decreases.

### Functional prediction

The abundance of each gene group has been normalized to the 16S rRNA copy number for each genome. The functional prediction results focus on eight distinct gene groups: Phototrophy, Lithotrophy, Organotrophy, DNA Conservation, DNA Repair, Nitrogen cycle, Sporulation and ROS-damage prevention (listed in Table S6). Figure 4 shows the pattern of the obtained functions. It shows higher abundances of the gene groups encoding for DNA conservation, DNA repair, nitrogen metabolism, ROS-damage prevention, sporulation, and phototrophy in patches associated with at least one EE compared to the barren patches (Table S7). Therefore, we analysed the results as barren vs average of the other three patch types that were not significantly different from one another (Table S8) and significant differences (p <0.04) between barren and EE(s) patches were detected. The genes related to lithotrophy, only differed between patches with one EE and the barren patches (p < 0.03) but patches with two EEs were similar to the barren plots. Finally, for genes related to the organotrophy, there was no significant differences between the patches (p>0.05).

**Figure 4.**
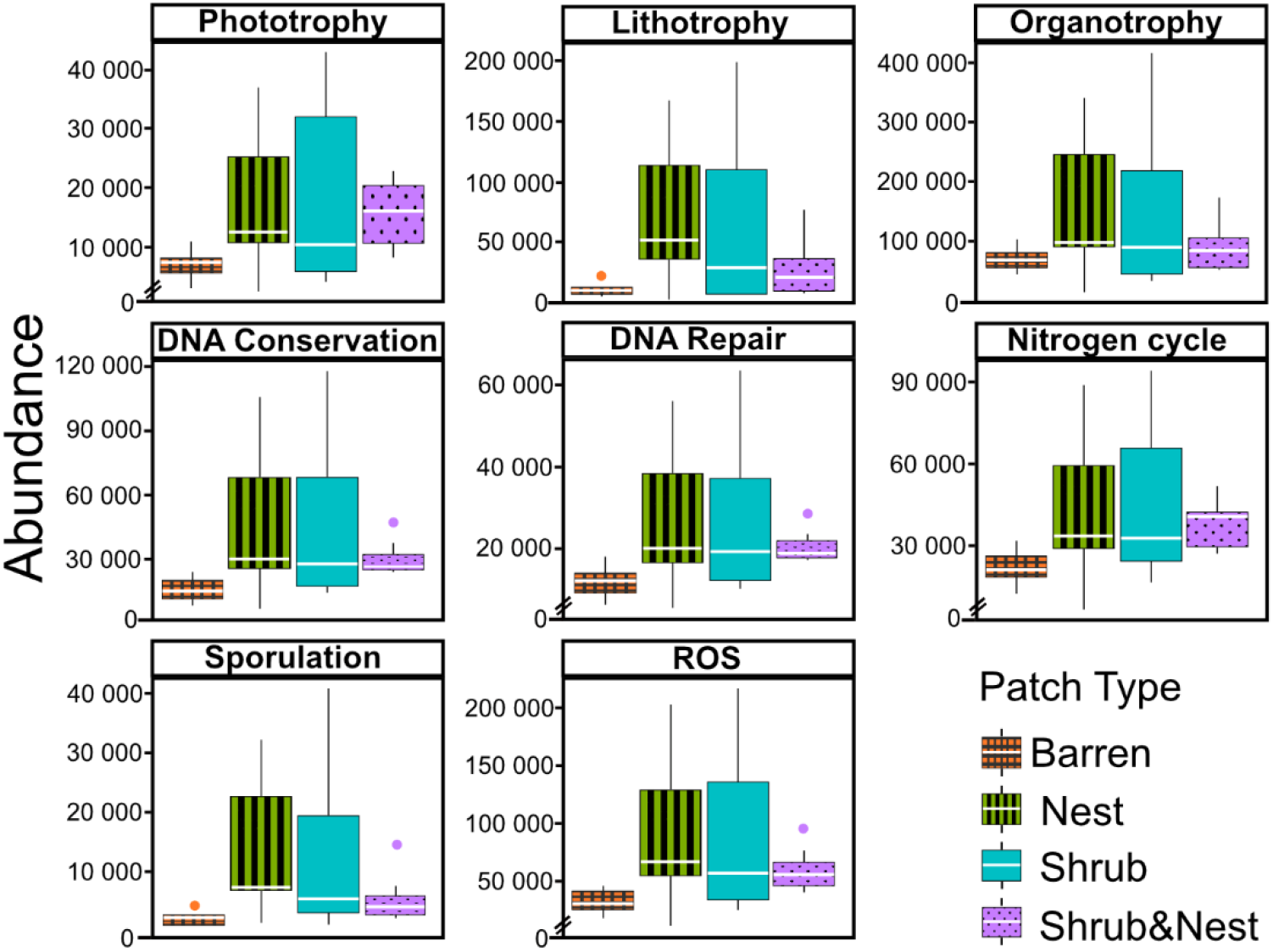
Boxplots of the functional prediction of the 16S sequences. Each panel (Boxplot) represents a different group of genes associated with a certain functionality. The full list of genes can be found in Table S4. The patch types are represented by distinct colours and patterns. The y-axis is the abundance in copy number (CN) normalized to the 16S rRNA copy number for each genome.

## DISCUSSION

In desert environments, during the dry season, a large portion of the microbial community is dormant or showing reduced metabolic activity (45, 58–60). However, the presence of EEs enhances the metabolic potential for metabolism-related and the survival-related functions (Figure 4). This may imply that the soil microbial communities occupying EE patches are better adapted to confront stressful events (e.g. rain event or desiccation). In addition, these communities may experience less extreme conditions that elsewhere due to the modulating effects of the EEs of the environmental conditions. Nevertheless, both the metabolism-related and the survival-related functions are enhanced by the presence of the various EEs evaluated here. The increase in the activity of the gene groups can be explained by an increase in nutrients in the combined EEs patches. However the physico-chemical measures, including soil water content, organic matter, nitrogen, phosphorus, and pH, did not match the changes observed in bacterial composition or function (Table S4, S5 and Figure 1) as was previously reported (13, 61, 62). We have previously proposed that the observed differences in community could be mediated by microclimatic characteristics under shrub patches (13). It has been reported that the desert dwarf shrubs affect the physical features of their immediate soil patch. Shrubs were shown to divert water flow and reduce evapotranspiration rates following rain events (63–65) and reduce temperature and radiation year round (66). Likewise, ants aerate the soil thus increasing infiltration during rain events (67) and mix the layers through bioturbation (18). We thus hypothesize that the prolonged water availability and altered physical conditions from the wet season shaped the community (68), establishing the composition and function observed here (Figure 3 and 4).

Two genera dominated the bacterial communities, both well adapted to stress conditions: *Rubrobacter* dominated the barren soil, while and *Deinococcus* dominated the EE patches (Figure 3 and Table S2). *Rubrobacter* are specialized in surviving strong desiccation and low nutrients (69, 70) showing high relative abundance in arid barren soils of the Negev highlands. *Deinococcus* are highly adapted to a wide range of extremes, such radiations, temperatures and, xerification. Some of these extreme conditions occur in the desert, while others are found in different environment, making *Deinococcus-Thermus* versatile organisms (57, 71, 72). We thus suggest that EE patches select for *Deinococcus* as they might be broader spectrum and can better adapt to perturbations compared to *Rubrobacter*

### The non-additive effects of two joint EEs

Only the combination of EEs resulted in significant changes (p-values: Table S5) of Nitrate, Phosphorus, and, to a lesser extent, Ammonium, pH, and OM (variables: Table S4). When located under a shrub, ants can increase their seed consumption, which enhances the amount of leftovers around the nest (73), and may increase the concentrations of nitrate and phosphorus. These macronutrients are important drivers of the biological processes, as they are often the limiting factors of microbial growth and activity in the soil (49). The EE patches analysed in this study share the same habitat and resources but their impact is separated, thus their joint impact is non-additive (74). The impact of an EE is defined by its lifetime, its population density, its spatial distribution, the time period of its presence on the site, the durability of its impact in the absence of other EEs, and the number, type, and magnitude of resource flows that are modified (8)(8). The behaviour of each EE is important as it becomes a feature of the combined impact of both EEs (75). However, the effect of both EEs together cannot be inferred from their individual environmental impact or from their mutual interaction (Figure 1) (30). Here, we investigated a sessile organism with a passive and slow impact (the perennial shrub) and compared it to a motile organism (the ants) with an active and impact that has both a short-term component through the seasonal accumulation of seeds and organic matter and a lasting impact due to the alternation of the nest mound which remains in the same place for decades (76). Even though, the impact of these EEs can be affiliated to the concept of resource islands (21), by creating havens of resources and water, their individual, and combined, effects do not always lead to strong changes in the composition of the soil microbial community (Figure 3). However, they create favorable conditions increasing the activity of the subsoil bacterial community (Figure 4) (13).

In our ecosystem, shrubs and ants are not the only two EEs and further studies should also consider the impact of other EEs. For example, the soil crust and the cyanobacteria living in it are recognized as important ecosystem engineers in arid ecosystems (8, 30, 77, 78). Furthermore, the soil crust in our system is often disturbed by the action of the other two EEs (6, 79). Thus, this third type of EE is not only important for its potential impact on microbial community composition and soil physico-chemical properties (80), but its distribution is also not independent of those of the other two EEs. Such complicated relationships may explain some of the discrepancies in our study.

## ACKOWLEDGMENTS

This study was supported by the Koshland Foundation to Itamar Giladi and Osnat Gillor.

## REFERENCES

1. Ben-David EA, Zaady E, Sher Y, Nejidat A. 2011. Assessment of the spatial distribution of soil microbial communities in patchy arid and semi-arid landscapes of the Negev Desert using combined PLFA and DGGE analyses. FEMS Microbiol Ecol 76:492–503.

2. Schlesinger WH, Raikks JA, Hartley AE, Cross AF. 1996. On the spatial pattern of soil nutrients in desert ecosystems. Ecology 77:364–374.

3. West NE. 1981. Nutrient cycling in desert ecosystems, p. 301–324. In Arid land ecosystems: Volume 2, Structure, functioning and management.

4. Wilby A, Shachak M, Boeken B. 2001. Integration of ecosystem engineering and trophic effects of herbivores. Oikos 92:436–444.

5. Wright JP, Jones CG, Boeken B, Shachak M. 2006. Predictability of ecosystem engineering effects on species richness across environmental variability and spatial scales: *Shrub mound effects on annual plant diversity*. J Ecol 94:815–824.

6. Oren Y, Perevolotsky A, Brand S, Shachak M. 2007. Livestock and engineering network in the Israeli Negev: Implications for ecosystem management, p. 323–342. In Ecosystem Engineers. Elsevier Inc.

7. Gosselin EN, Holbrook JD, Huggler K, Brown E, Vierling KT, Arkle RS, Pilliod DS. 2016. Ecosystem engineering of harvester ants: effects on vegetation in a sagebrush-steppe ecosystem. West North Am Nat 76:82–89.

8. Jones CG, Lawton JH, Shachak M. 1994. Organisms as Ecosystem Engineers. Oikos 69:373–386.

9. Lavelle P. 2002. Functional domains in soils. Ecol Res 17:441–450.

10. Lavelle P, Blanchart E, Martin A, Spain A V., Martin S. 1992. Impact of Soil Fauna on the Properties of Soils in the Humid Tropics. Soil Sci Soc Am https://doi.org/10.2136/sssaspecpub29.c9.

11. Lavelle P, Decaëns T, Aubert M, Barot S, Blouin M, Bureau F, Margerie P, Mora P, Rossi JP. 2006. Soil invertebrates and ecosystem services. Eur J Soil Biol 42.

12. Wright SF, Upadhyaya A. 1996. Extraction of an abundant and unusual protein from soil and comparison with hyphal protein of arbuscular mycorrhizal fungi. Soil Sci 161.

13. Bachar A, Soares MIM, Gillor O. 2012. The effect of resource islands on abundance and diversity of bacteria in arid soils. Microb Ecol 63:694–700.

14. Ginzburg O, Whitford WG, Steinberger Y. 2008. Effects of harvester ant (Messor spp.) activity on soil properties and microbial communities in a Negev Desert ecosystem. Biol Fertil Soils 45:165–173.

15. Saul-Tcherkas V, Steinberger Y. 2011. Soil microbial diversity in the vicinity of a Negev desert shrub-Reaumuria negevensis. Microb Ecol 61:64–81.

16. Camargo-Ricalde SL, Dhillion SS. 2003. Endemic Mimosa species can serve as mycorrhizal “resource islands” within semiarid communities of the Tehuacán-Cuicatlán Valley, Mexico. Mycorrhiza 13:129–136.

17. Robinson BS, Bamforth SS, Dobson PJ. 2002. Density and diversity of protozoa in some arid Australian soils. J Eukaryot Microbiol 49:449–453.

18. Folgarait P. 1998. Ant biodiversity to ecosystem functioning: a review. Biodivers Conserv 7:1121–1244.

19. MacMahon JA, Mull JF, Crist TO. 2000. Harvester ants (Pogonomyrmex spp.): their community and ecosystem influences. Annu Rev Ecol Syst 31:265–291.

20. Callaway RM. 1995. Positive interactions among plants. Bot Rev 61:306–349.

21. Schlesinger WH, Pilmanis AM. 1998. Plant-soil Interactions in Deserts. Biogeochemistry 42:169–187.

22. Segoli M, Ungar ED, Giladi I, Arnon A, Shachak M. 2012. Untangling the positive and negative effects of shrubs on herbaceous vegetation in drylands. Landsc Ecol 27:899–910.

23. Shachak M, Boeken B, Groner E, Kadmon R, Lubin Y, Meron E, Ne’eman G, Perevolotsky A, Shkedy Y, Ungar ED. 2008. Woody species as landscape modulators and their effect on biodiversity patterns. Bioscience 58:209–221.

24. Walker LR, Thompson DB, Landau FH. 2001. Experimental manipulations of fertile islands and nurse plant effects in the Mojave Desert, USA. West North Am Nat 61:25–35.

25. Steven B, Gallegos-Graves LV, Yeager C, Belnap J, Kuske CR. 2014. Common and distinguishing features of the bacterial and fungal communities in biological soil crusts and shrub root zone soils. Soil Biol Biochem 69:302–312.

26. Facelli JM, Temby AM. 2002. Multiple effects of shrubs on annual plant communities in arid lands of South Australia. Austral Ecol 27:422–432.

27. Pariente S. 2002. Spatial patterns of soil moisture as affected by shrubs, in different climatic conditions. Environ Monit Assess 73:237–251.

28. Frouz J, Holec M, Kalčík J. 2003. The effect of Lasius niger (Hymenoptera, Formicidae) ant nest on selected soil chemical properties. Pedobiologia (Jena) 47:205–212.

29. Farji-Brener AG, Werenkraut V. 2017. The effects of ant nests on soil fertility and plant performance: a meta-analysis. J Anim Ecol 86:866–877.

30. Gilad E, von Hardenberg J, Provenzale A, Shachak M, Meron E. 2004. Ecosystem Engineers: From Pattern Formation to Habitat Creation. Phys Rev Lett 93:098105.

31. Angel R. 2012. Total Nucleic Acid Extraction from Soil.

32. Klindworth A, Pruesse E, Schweer T, Peplies J, Quast C, Horn M, Glöckner FO, Glockner FO. 2013. Evaluation of general 16S ribosomal RNA gene PCR primers for classical and next-generation sequencing-based diversity studies. Nucleic Acids Res 41:1–11.

33. SSSA SSS of A. 1996. Methods of Soil Analysis: Part 3 Chemical methods, 5.3.

34. R Core Team I. 2016. R: A language and environment for statistical computing. R Found Stat Comput.

35. Dinno A. 2017. Package ‘dunn.test.’ CRAN Repos 1–7.

36. Dunn OJ. 1964. Multiple Comparisons Using Rank Sums. Technometrics 6:241–252.

37. Kruskal WH, Wallis WA. 1952. Use of Ranks in One-Criterion Variance Analysis. J Am Stat Assoc 47:583–621.

38. Bolyen E, Rideout JR, Dillon MR, Bokulich NA, Abnet C, Al-Ghalith GA, Alexander H, Alm EJ, Arumugam M, Asnicar F, Bai Y, Bisanz JE, Bittinger K, Brejnrod A, Brislawn CJ, Brown CT, Callahan BJ, Caraballo-Rodríguez AM, Chase J, Cope E, Da Silva R, Dorrestein PC, Douglas GM, Durall DM, Duvallet C, Edwardson CF, Ernst M, Estaki M, Fouquier J, Gauglitz JM, Gibson DL, Gonzalez A, Gorlick K, Guo J, Hillmann B, Holmes S, Holste H, Huttenhower C, Huttley G, Janssen S, Jarmusch AK, Jiang L, Kaehler B, Kang K Bin, Keefe CR, Keim P, Kelley ST, Knights D, Koester I, Kosciolek T, Kreps J, Langille MGI, Lee J, Ley R, Liu Y-X, Loftfield E, Lozupone C, Maher M, Marotz C, Martin BD, McDonald D, McIver LJ, Melnik A V, Metcalf JL, Morgan SC, Morton J, Naimey AT, Navas-Molina JA, Nothias LF, Orchanian SB, Pearson T, Peoples SL, Petras D, Preuss ML, Pruesse E, Rasmussen LB, Rivers A, Robeson Michael S II, Rosenthal P, Segata N, Shaffer M, Shiffer A, Sinha R, Song SJ, Spear JR, Swafford AD, Thompson LR, Torres PJ, Trinh P, Tripathi A, Turnbaugh PJ, Ul-Hasan S, van der Hooft JJJ, Vargas F, Vázquez-Baeza Y, Vogtmann E, von Hippel M, Walters W, Wan Y, Wang M, Warren J, Weber KC, Williamson CHD, Willis AD, Xu ZZ, Zaneveld JR, Zhang Y, Zhu Q, Knight R, Caporaso JG. 2018. QIIME 2: Reproducible, interactive, scalable, and extensible microbiome data science. PeerJ Prepr 6:e27295v2.

39. Callahan BJ, McMurdie PJ, Rosen MJ, Han AW, Johnson AJA, Holmes SP. 2016. DADA2: High-resolution sample inference from Illumina amplicon data. Nat Methods2016/05/23. 13:581–583.

40. Quast C, Pruesse E, Yilmaz P, Gerken J, Schweer T, Yarza P, Peplies J, Glöckner FO. 2013. The SILVA ribosomal RNA gene database project: improved data processing and web-based tools. Nucleic Acids Res 41:D590–D596.

41. Oksanen J, Blanchet FG, Kindt R, Legen-P, Minchin PR, Hara RBO, Simpson GL, Solymos P, Stevens MHH. 2014. Package ‘ vegan ’ https://doi.org/ISBN0-387-95457-0.

42. Narayan NR, Weinmaier T, Laserna-Mendieta EJ, Claesson MJ, Shanahan F, Dabbagh K, Iwai S, Desantis TZ. 2020. Piphillin predicts metagenomic composition and dynamics from DADA2-corrected 16S rDNA sequences. BMC Genomics 21:1–12.

43. Iwai S, Weinmaier T, Schmidt BL, Albertson DG, Poloso NJ, Dabbagh K, DeSantis TZ. 2016. Piphillin: Improved prediction of metagenomic content by direct inference from human microbiomes. PLoS One 11:1–18.

44. Kaneshisa M, Goto S. 2000. KEGG: Kyoto Encyclopedia of Genes and Genomes. Nucleic Acids Res 28:27–30.

45. Cordero PRF, Bayly K, Man Leung P, Huang C, Islam ZF, Schittenhelm RB, King GM, Greening C. 2019. Atmospheric carbon monoxide oxidation is a widespread mechanism supporting microbial survival. ISME J 13:2868–2881.

46. Greening C, Biswas A, Carere CR, Jackson CJ, Taylor MC, Stott MB, Cook GM, Morales SE. 2016. Genomic and metagenomic surveys of hydrogenase distribution indicate H 2 is a widely utilised energy source for microbial growth and survival. ISME J 10:761–777.

47. León-Sobrino C, Ramond JB, Maggs-Kölling G, Cowan DA. 2019. Nutrient acquisition, rather than stress response over diel cycles, drives microbial transcription in a hyper-arid Namib desert soil. Front Microbiol 10:1–11.

48. Tveit AT, Hestnes AG, Robinson SL, Schintlmeister A, Dedysh SN, Jehmlich N, Von Bergen M, Herbold C, Wagner M, Richter A, Svenning MM. 2019. Widespread soil bacterium that oxidizes atmospheric methane. Proc Natl Acad Sci U S A 116:8515–8524.

49. Madigan MT, Martinko JM, Dunlap P V, Clark DP. 2009. Brock Biology of microoragnisms.

50. Borisov VB, Forte E, Davletshin A, Mastronicola D, Sarti P, Giuffrè A. 2013. Cytochrome bd oxidase from Escherichia coli displays high catalase activity: An additional defense against oxidative stress. FEBS Lett 587:2214–2218.

51. Hansen BB, Henriksen S, Aanes R, Sæther BE. 2007. Ungulate impact on vegetation in a two-level trophic system. Polar Biol 30:549–558.

52. Henrikus SS, Wood EA, McDonald JP, Cox MM, Woodgate R, Goodman MF, van Oijen AM, Robinson A. 2018. DNA polymerase IV primarily operates outside of DNA replication forks in Escherichia coli. PLoS Genet 14:1–29.

53. Preiss J. 1984. Bacterial glycogen synthesis and its regulation. Annu Rev Microbiol 38:419–458.

54. Preiss J, Sivak M. 1999. 3.14-Starch and Glycogen Biosynthesis, p. 441–495. In Barton, SD, Nakanishi, K, Meth-Cohn, OBT-CNPC (eds.),. Pergamon, Oxford.

55. Rajeev L, Da Rocha UN, Klitgord N, Luning EG, Fortney J, Axen SD, Shih PM, Bouskill NJ, Bowen BP, Kerfeld CA, Garcia-Pichel F, Brodie EL, Northen TR, Mukhopadhyay A. 2013. Dynamic cyanobacterial response to hydration and dehydration in a desert biological soil crust. ISME J 7:2178–2191.

56. Repar J, Briski N, Buljubašić M, Zahradka K, Zahradka D. 2012. Exonuclease VII is involved in “reckless” DNA degradation in UV-irradiated Escherichia coli. Mutat Res 750.

57. Slade D, Radman M. 2011. Oxidative Stress Resistance in Deinococcus radioduransMicrobiology and Molecular Biology Reviews.

58. Bay S, Ferrari B, Greening C. 2018. Life without water: How do bacteria generate biomass in desert ecosystems? Microbiol Aust 39:28–32.

59. Lennon JT, Jones SE. 2011. Microbial seed banks: the ecological and evolutionary implications of dormancy. Nat Rev Micro 9:119–130.

60. Schulze-Makuch D, Wagner D, Kounaves SP, Mangelsdorf K, Devine KG, de Vera J-P, Schmitt-Kopplin P, Grossart H-P, Parro V, Kaupenjohann M, Galy A, Schneider B, Airo A, Frösler J, Davila AF, Arens FL, Cáceres L, Cornejo FS, Carrizo D, Dartnell L, DiRuggiero J, Flury M, Ganzert L, Gessner MO, Grathwohl P, Guan L, Heinz J, Hess M, Keppler F, Maus D, McKay CP, Meckenstock RU, Montgomery W, Oberlin EA, Probst AJ, Sáenz JS, Sattler T, Schirmack J, Sephton MA, Schloter M, Uhl J, Valenzuela B, Vestergaard G, Wörmer L, Zamorano P. 2018. Transitory microbial habitat in the hyperarid Atacama Desert. Proc Natl Acad Sci 115:2670–2675.

61. Angel R, Soares MIM, Ungar ED, Gillor O. 2010. Biogeography of soil archaea and bacteria along a steep precipitation gradient. ISME J 4:553–563.

62. Vonshak A, Sklarz MY, Hirsch AM, Gillor O. 2018. Perennials but not slope aspect affect the diversity of soil bacterial communities in the northern Negev Desert, Israel. Soil Res 56:123–128.

63. Sarig S, Steinberger Y. 1993. Microbial pools of N and P under desert halophyte shrubs: Response to soil salinity. Beyond biomass Compos Funct Anal soil Microb communities.

64. Whitford WG, Duval BD. 2002. Ecology of Desert SystemsAgriculture, Ecosystems & Environment.

65. Segoli M, Ungar ED, Shachak M. 2008. Shrubs enhance resilience of a semi-arid ecosystem by engineering and regrowth. Ecohydrology 1:330–339.

66. Kidron GJ. 2009. The effect of shrub canopy upon surface temperatures and evaporation in the Negev Desert. Earth Surf Process Landforms 34:123–132.

67. Berg N, Steinberger Y. 2008. Role of perennial plants in determining the activity of the microbial community in the Negev Desert ecosystem. Soil Biol Biochem https://doi.org/10.1016/j.soilbio.2008.07.019.

68. Baubin C, Farrell AM, Šťovíček A, Ghazaryan L, Giladi I, Gillor O. 2019. Seasonal and spatial variability in total and active bacterial communities from desert soil. Pedobiologia (Jena) 74:7–14.

69. Bull AT. 2011. Actinobacteria of the Extremobiosphere, p. 1203–1240. In Horikoshi, K (ed.), Extremophiles Handbook. Springer Japan, Tokyo.

70. Ferreira AC, Nobre MF, Moore E, Rainey FA, Battista JR, Da Costa MS. 1999. Characterization and radiation resistance of new isolates of Rubrobacter radiotolerans and Rubrobacter xylanophilus. Extremophiles 3:235–238.

71. Chanal A, Chapon V, Benzerara K, Barakat M, Christen R, Achouak W, Barras F, Heulin T. 2006. The desert of Tataouine: An extreme environment that hosts a wide diversity of microorganisms and radiotolerant bacteria. Environ Microbiol 8:514–525.

72. Prieur D. 2007. An Extreme Environment on Earth: Deep-Sea Hydrothermal Vents. Lessons for Exploration of Mars and Europa, p. 319–345. In Gargaud, M, Martin, H, Claeys, P (eds.), Lectures in Astrobiology: Volume II. Springer Berlin Heidelberg, Berlin, Heidelberg.

73. Wagner D. 1997. The Influence of Ant Nests on Acacia Seed Production, Herbivory and Soil Nutrients. J Ecol 85:83–93.

74. Passarelli C, Olivier F, Paterson DM, Meziane T, Hubas C. 2014. Organisms as cooperative ecosystem engineers in intertidal flats. J Sea Res 92:92–101.

75. Alba-Lynn C, Detling JK. 2008. Interactive disturbance effects of two disparate ecosystem engineers in North American shortgrass steppe. Oecologia 157:269–278.

76. Wagner D, Jones JB. 2004. The Contribution of Harvester Ant Nests, Pogonomyrmex rugosus (Hymenoptera, Formicidae), to Soil Nutrient Stocks and Microbial Biomass in the Mojave Desert. Environ Entomol 33:599–607.

77. West NE. 1990. Structure and function of microphytic soil crusts in wildland ecosystems of arid to semi-arid regions. Adv Ecol Res https://doi.org/10.1016/S0065-2504(08)60055-0.

78. Eldridge DJ, Bowker MA, Maestre FT, Alonso P, Mau RL, Papadopoulos J, Escudero A. 2010. Interactive effects of three ecosystem engineers on infiltration in a semi-arid Mediterranean grassland. Ecosystems 13:499–510.

79. Li XR, Gao YH, Su JQ, Jia RL, Zhang ZS. 2014. Ants mediate soil water in arid desert ecosystems: Mitigating rainfall interception induced by biological soil crusts? Appl Soil Ecol 78:57–64.

80. Schulz K, Mikhailyuk T, Dreßler M, Leinweber P, Karsten U. 2016. Biological Soil Crusts from Coastal Dunes at the Baltic Sea: Cyanobacterial and Algal Biodiversity and Related Soil Properties. Microb Ecol 71:178–193.

